# Genetics of resistance to common root rot (spot blotch), *Fusarium* crown rot, and sharp eyespot in wheat

**DOI:** 10.1101/2020.07.30.228932

**Authors:** Jun Su, Jiaojie Zhao, Shuqing Zhao, Mengyu Li, Shuyong Pang, Shisheng Chen, Feng Chen, Wenchao Zhen, Zhensheng Kang, Xiaodong Wang

## Abstract

Due to soil changes, high density planting, and the use of straw-returning methods, wheat common root rot (spot blotch), *Fusarium* crown rot (FCR), and sharp eyespot (sheath blight) have become severe threats to global wheat production. Only a few wheat genotypes show moderate resistance to these root and crown rot fungal diseases, and the genetic determinants of wheat resistance to these devastating diseases are poorly understood. This review summarizes recent results of genetic studies of wheat resistance to common root rot, *Fusarium* crown rot, and sharp eyespot. Wheat germplasm with relatively higher resistance are highlighted and genetic loci controlling the resistance to each disease are summarized.

## Introduction

Long-term environmental changes have greatly affected crop diseases. For example, the higher temperatures associated with global warming may increase the severity of many plant diseases (Cohen and Leach 2020). Bursts of wheat stem base rot diseases, including common root rot, *Fusarium* crown rot, and sharp eyespot, are highly correlated with crop rotation practices. The large-scale application of wheat-maize rotation in the North China wheat cultivation area has dramatically changed the organic carbon, fertilization state, and nitrogen balance of the soil (Wang et al. 2015; Zhao et al. 2006). The disease suppressive capacity of the soil microbiome is also highly dependent on crop rotational diversity (Peralta et al. 2018).

### Pathogenic profiles

Wheat common root rot is caused by *Bipolaris sorokiniana* infection (**Fig. 1a**, teleomorph *Cochliobolus sativus*) in the root and stem base of wheat plants. Severe infections of this fungal pathogen in the root and crown of seedlings may kill plants. *B. sorokiniana* can also induce phenotypes of leaf spot (spot blotch, *Helminthosporium* leaf blight, or foliar blight, **Fig. 1b**), seedling wilt, head blight, and black point in *Triticeae* crops (Kumar et al. 2002). The average yield loss caused by *B. sorokiniana* ranges from 15% to 20%, but under favorable heat and drought conditions this disease can decrease wheat production by 70% and reduce seed quality (Sharma and Duveiller 2007). This fungal pathogen accumulates several toxins to kill or weaken plant cells, including prehelminthosporol, helminthosporol, helminthosporic acid, sorokinianin, and bipolaroxin (Kumar et al. 2002; Gupta et al. 2018). However, the potential negative effects of *B. sorokiniana*-infected wheat grains (black point) on food safety have not been investigated in detail. *B. sorokiniana* has a very wide host range, as it can infect wheat, barley, maize, rice, and many other grass species (Gupta et al. 2018). Multiple-year *Triticeae* crop rotations of wheat and barley greatly promote the severity of common root rot caused by *B. sorokiniana* (Conner et al. 1996). Maize crops and returned straws may also be infected by this fungus, so common root rot and spot blotch have been more frequently observed in areas of wheat cultivation in North China where methods of large-scale wheat-maize rotation and straw returning have been applied.

**Fig. 1.**
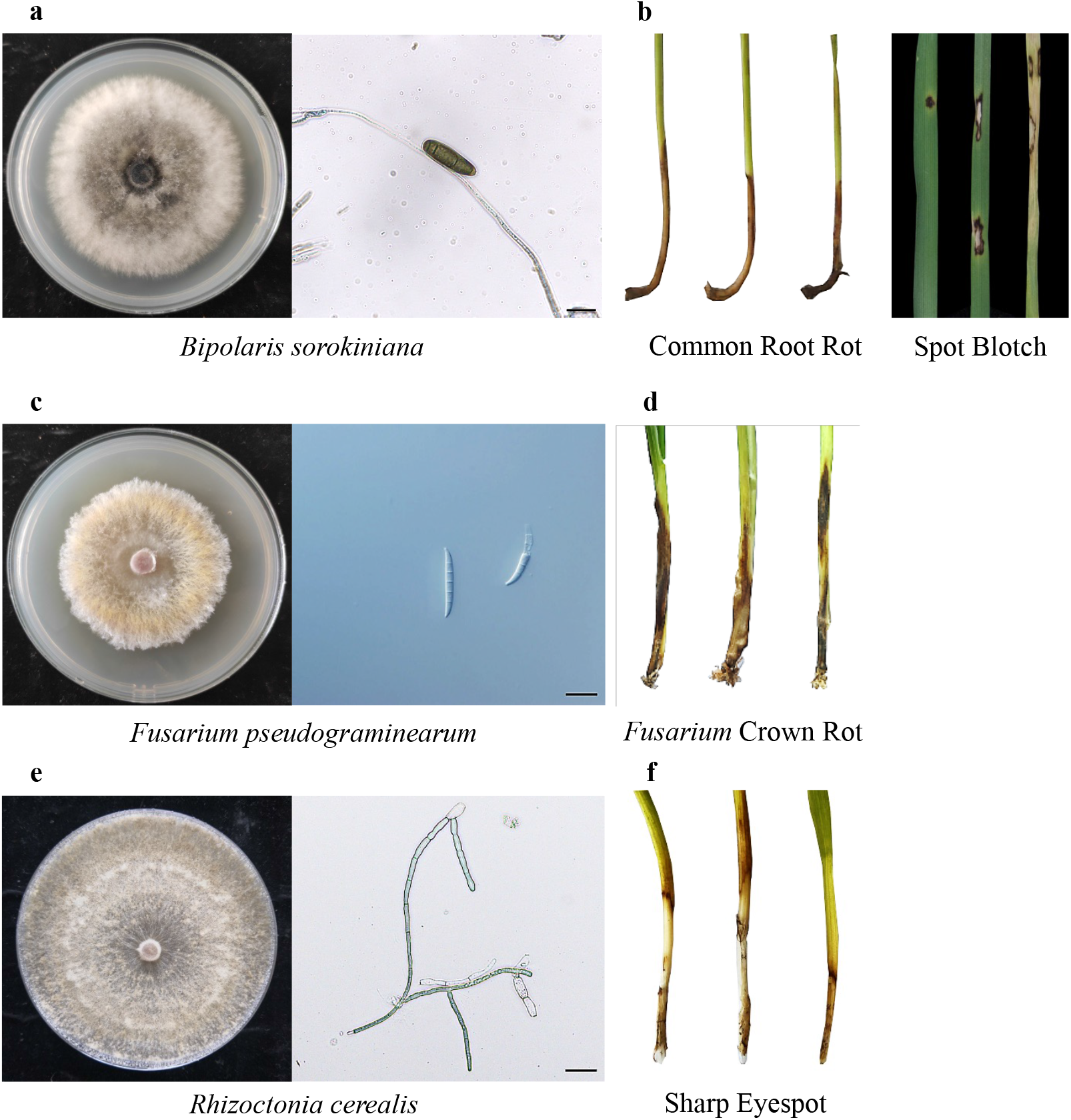
Pathogenic profiles of *Bipolaris sorokiniana, Fusarium pseudograminearum*, and *Rhizoctonia cerealis*. **a** *B. sorokiniana* was cultivated on potato dextrose agar (PDA) medium and spores were directly collected. **b** Common root rot and spot blotch caused by *B. sorokiniana*. Infected wheat plants were easily pulled out, the stem base and root system felt wet, and black and brown striped spots can be observed in both the stem base and lower leaves. **c** *F. pseudograminearum* cultivated on PDA medium. Spores of *F. pseudograminearum* can be induced on carboxymethyl cellulose sodium (CMC) medium. **d** *Fusarium* crown rot caused by *F. pseudograminearum*. The stem base of infected wheat plants became dry and fragile, and was easily broken apart. Additionally, dark and red brown rot can be observed in the stem base. **e** *R. cerealis* was cultivated on PDA medium. **f** Sharp eyespot caused by *R. cerealis*. The typical lesions on wheat stem are elliptical or exhibit an “eye” shape with sharply dark brown borders. Scale bar = 20 µm.

*Fusarium* crown rot (FCR) is caused by infection of *Fusarium pseudograminearum* (**Fig. 1c**), or other *Fusarium* pathogens including *F. culmorum, F. avenaceum*, and *F. graminearum*. These fungal species infect the coleoptile, leaf sheath, and stem base of wheat seedlings, generating browning and decay phenotypes (**Fig. 1d**). *Fusarium* pathogens are found globally in arid and semi-arid wheat planting areas (Kazan and Gardiner 2018). FCR infection caused an estimated 35% yield loss of winter wheat in the Northwest Pacific region of the United States (Smiley et al. 2005). When FCR-infected plants are co-infected with *Fusarium* Head Blight (FHB), wheat seeds are likely to be contaminated by fungal toxins such as deoxynivalenol (DON) and nivalenol (NIV), which greatly threaten the health of human and livestock (Obanor and Chakraborty 2014; Monds et al. 2005). Maize also can be infected with various *Fusarium* pathogens, and the fungi from infected plants can remain active in returned straw debris for as long as five years (Burgess et al. 2001). For these reasons, FCR is a growing threat to wheat cultivation in wheat-maize rotation regions in North China.

Wheat sharp eyespot (sheath blight) is caused by infection of *Rhizoctonia cerealis* (**Fig. 1e**) in the root and stem base of wheat plants, generating disease phenotypes of stem eyespot (**Fig. 1f**), crown rot, seedling fatal damage, and head blight. Wheat sharp eyespot is a typical soil-borne fungal disease that is prevalent worldwide (Hamada et al. 2011). *R. cerealis* also has a broad host range, including many cereals. This fungal pathogen can survive in soil or on infected crop residues for a long time. Consequently, practices of wheat-maize rotation and straw-returning have greatly facilitated the burst of this disease in China during the last two decades (Ren et al. 2020). In 2005, approximately 8 million ha of wheat fields in China were infected with sharp eyespot, with an estimated yield loss of about 530,000 tons (McBeath and McBeath 2010). Sharp eyespot also significantly decreases wheat grain quality (Lemańczyk and Kwaśna 2013).

These three diseases can have similar phenotypes, causing stem base rot and head blight, but there are differences as well. Common root rot caused by *B. sorokiniana* weakens infected wheat plants so they can be easily pulled out. Additionally the stem base and root system feel wet, and black and brown striped spots form on both the stem base and lower leaves (**Fig. 1b**). For FCR caused by *F. pseudograminearum*, the stem base of the infected wheat plant becomes dry and fragile, and dark brown rot can be observed in the stem base (**Fig. 1d**). For sharp eyespot caused by *R. cerealis*, most lesions on the wheat stem are elliptical or have a “eye” shape, with sharply dark brown borders (**Fig. 1f**).

### Progress in dissecting the genetic determinants of wheat resistance to common root rot (spot blotch)

The use of wheat resistant cultivars remains the most efficient and economical way to control common root rot (spot blotch). However, there are currently insufficient germplasm resources with resistance to common root rot to meet the growing needs for global wheat breeding applications and there have been few studies to identify the genetic loci that control resistance to common root rot (Gupta et al. 2018). Early efforts focused on the introgression of common root rot resistant loci from *Thinopyrum ponticum*, a wheat relative (Li et al. 2004). Wheat breeding programs for common root rot resistance have had limited success because analysis of complex quantitative trait loci (QTL) is required (Joshi et al. 2004). Using bi-parental populations and linkage mapping, four genetic loci with major resistant effect were identified and designated as *Sb* genes. *Sb1* was discovered in the bread wheat line “Saar”, was mapped to chromosome 7DS, and is associated with the wheat leaf rust resistance gene *Lr34* (Lillemo et al. 2013). The *Lr34/Yr18/Pm38* gene encodes a ATP-binding cassette (ABC) transporter that confers broad-spectrum resistance to multiple foliar fungal diseases, including leaf rust, stripe rust, and powdery mildew (Krattinger et al. 2009). Another minor QTL linked to *Lr46* on chromosome 1BL was also identified from “Saar”. The *Lr46* gene is associated with resistance to leaf rust in adult plants and is also associated with the stripe rust resistance gene *Yr29* (William et al. 2003). The *Sb2* gene was identified in bread wheat cultivar “Yangmai 6” and was mapped to chromosome 5BL between simple sequence repeat (SSR) markers of *Xgwm639* and *Xgwm1043* (Kumar et al. 2015). The *Sb2* gene was later reported to be linked with the *Tsn1* gene, which confers host-selective sensitivity to the fungal toxin ToxA produced by *Pyrenophora tritici-repentis* (Kumar et al. 2016). The *Sb3* gene was discovered in the winter wheat line “621-7-1” based on its correlation with immune response to *B. sorokiniana*. Using bulked segregant analysis (BSA), *Sb3* was mapped to chromosome 3BS, flanking SSR markers of *Xbarc133* and *Xbarc147* (Lu et al. 2016). The *Sb4* gene was recently identified from two highly resistant wheat lines, “Zhongyu1211” and “GY17”. Using RNA-based BSA and single-nucleotide polymorphism (SNP) mapping, *Sb4* was delimitated to a 1.19 cM genetic interval region of chromosome 4BL, which contains 21 predicted genes in the corresponding “Chinese Spring” genome (Zhang et al. 2020). Future work should clone these *Sb* genes to further elucidate the mechanism of wheat resistance toward this devastating fungal pathogen.

Several other major QTLs have been discovered and preliminarily mapped using bi-parental populations. For example, two resistant QTLs derived from “Yangmai 6” were mapped to chromosomes 5B and 7D using microsatellite markers (Kumar et al. 2005). Three QTLs on chromosomes 5B, 6A, and 6D were identified based on analysis of SSR markers from the resistant genotype “G162” (Sharma et al. 2007). Four QTLs controlling resistance of wheat cultivar “Yangmai 6” to *B. sorokiniana* were mapped to chromosomes 2AL, 2BS, 5BL, and 6DL (Kumar et al. 2009). A total of seven QTLs providing resistance to *B. sorokiniana* infections were mapped in the wheat lines “Ning 8201” and “Chirya 3” (Kumar et al. 2010). Three QTLs on chromosomes 1BS, 3BS, and 5AS respectively explained 8.5%, 17.6%, and 12.3%, of the resistant effect in “SYN1”, a CIMMYT (International Maize and Wheat Improvement Center) synthetic-derived bread wheat line (Zhu et al. 2014). From the Brazilian resistant cultivar “BH 1146”, two QTLs on chromosomes 7BL and 7DL were mapped using microsatellite markers (Singh et al. 2016). A prominent resistant QTL near the *Vrn-A1* locus on chromosome 5AL was also found in “BARTAI” and “WUYA” CIMMYT breeding lines (Singh et al. 2018). Finally, QTLs in *Vrn-A1* and *Sb2/Tsn1* loci were detected in two other CIMMYT breeding lines, “CASCABEL” and “KATH” (He et al.).

Genome-wide association studies (GWAS) have been widely used to identify QTLs. Using 832 polymorphic Diversity Arrays Technology (DArT) markers, four QTLs resistant to spot blotch were mapped to chromosomes 1A, 3B, 7B, and 7D after analysis of 566 spring wheat germplasm (Adhikari et al. 2012). A phenotypic screening of 11 parental genotypes and 55 F_2_ lines identified “19HRWSN6” as a resistant source. Subsequent simple linear regression analysis revealed SSR markers on chromosomes 5B, 6A, and 7D associated with resistance to *B. sorokiniana* (Tembo et al. 2017). There has been recent progress in drafting the physical genome of hexaploid wheat (Appels et al. 2018), and high-throughput SNP toolkits are now available for GWAS on various complex traits of wheat (Sun et al. 2020). A total of 528 spring wheat genotypes from different geographic regions were tested for spot blotch resistance and eleven associated SNP markers were found by 9K SNP assay (Gurung et al. 2014). Another study evaluated the responses of 294 hard winter wheat genotypes to *B. sorokiniana* and performed GWAS by 15K SNP assay. Ten wheat genotypes with relatively high resistance were identified, and six major resistant QTLs were found to collectively explain 30% of the phenotypic variation (Ayana et al. 2018). A total of 159 spring wheat genotypes were screened for common root rot resistance and twenty-four QTLs were identified, with a major one on chromosome 7B that explained 14% of the phenotypic variation of spot blotch severity (Jamil et al. 2018). Another study profiled the resistant phenotype of 287 spring wheat germplasm and performed GWAS using 90K SNP array. Eight genetic loci were associated with incubation period, lesion number, and disease score of *B. sorokiniana* infection (Ahirwar et al. 2018). A recent study phenotyped 301 Afghan wheat germplasm and found that approximately 15% exhibited lower disease scores than the resistant control. A subsequent GWAS approach identified twenty-five marker-trait associations on more than twelve chromosomes, including previously identified *Vrn-A1* and *Sb2/Tsn1* loci (Bainsla et al.). Another 141 spring wheat lines were collected for GWAS on spot blotch resistance. A total of 23 genomic regions were identified, including several stable regions on chromosomes 2B, 5B and 7D, and a novel region on chromosome 3D (Tomar et al. 2020).

We have summarized the previously reported wheat germplasm with relatively higher resistance to *B. sorokiniana* **(Supplementary Table 1)**. These wheat materials may serve as valuable resources for the genetic improvement of wheat resistance to common root rot (spot blotch). We have also summarized detailed information of previously designated resistant QTLs **(Supplementary Table 1)** and drafted their genomic distributions using the released genome of hexaploid wheat **(Fig. 2)**.

**Fig. 2.**
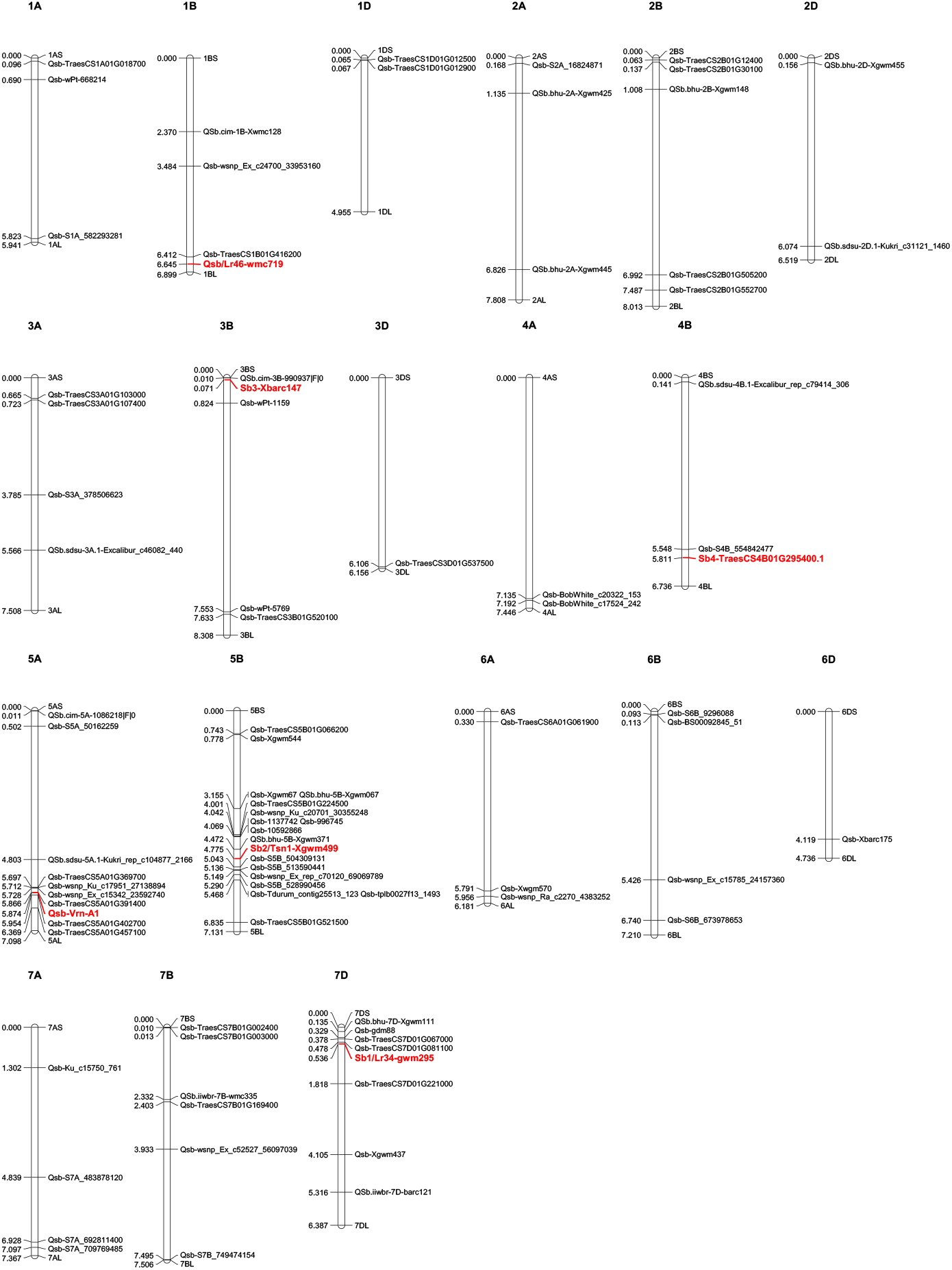
Genetics of resistance to common root rot (spot blotch) in wheat. Molecular markers, SNPs, and genes associated with common root rot or spot blotch resistant QTLs were collected from previous publications and searched against the JBrowse-1.12.3-release of the common wheat “Chinese Spring” genome available from the “*Triticeae* Multi-omics Center (http://202.194.139.32/)”. Physical positions (numbers indicated on the left side of each chromosome) were used to generate a distribution map of all the collected QTLs using Mapchart v2.32 software. QTLs with major effect or linked with designated genes are highlighted in red. Detailed information for these QTLs can be found in **Supplementary Table 1**.

### Genetic loci controlling wheat resistance to *Fusarium* crown rot

Since the causal agent of *Fusarim* head blight (FHB), *Fusarium graminearum*, can also induce the phenotype of *Fusarium* crown rot (Zhou et al. 2019; Akinsanmi et al. 2006), it is likely that FHB-resistant germplasm and genetic loci can be exploited to improve FCR resistance. For instance, the recently cloned FHB resistance gene *Fhb7* encodes a glutathione S-transferase (GST) and provides broad resistance to *Fusarium* diseases, including FCR induced by *F. pseudograminearum*, by detoxifying trichothecenes through de-epoxidation (Wang et al. 2020). However, an earlier investigation of the same wheat genotypes found no significant correlation of resistant phenotype or genetic loci conferring resistance to FHB and FCR (Li et al. 2010). A recent large-scale phenotyping of 205 Chinese wheat cultivars for resistance to both FHB and FCR also found no correlation in resistant phenotypes (Shi et al. 2020). Great efforts have also been made towards identification of FCR-resistant barley germplasm and genetic loci that control FCR resistance in barley (Liu and Ogbonnaya 2015). Since recent review papers have already summarized QTLs conferring FHB resistance and susceptibility in wheat in detail (Buerstmayr et al. 2020; Fabre et al. 2020), here we have mainly focused on studies reporting wheat resistance to FCR induced by *F. pseudograminearum* and *F. culmorum*.

Genetic studies revealed a major FCR-resistant QTL on chromosome 3BL (*Qcrs*.*cpi-3B*). This resistant locus, *Qcrs*.*cpi-3B*, was identified in the wheat genotype “CSCR6” of the taxon *Triticum spelta* (Ma et al. 2010). In a wheat recombinant inbred line population of “Lang/CSCR6”, a QTL on chromosome 4B derived from “Lang” explained the soil-free FCR resistance (Yang et al. 2010). Another significant QTL on chromosome 6B was identified as responsible for FCR resistance during an introgression process for durum wheat using “CSCR6” as the donor parent (Ma et al. 2012b). Near-isogenic lines for the *Qcrs*.*cpi-3B* locus have been developed for both genetic research and breeding practice (Ma et al. 2012a). Subsequent transcriptome and allele specificity analysis revealed differentially expressed genes associated with the *Qcrs*.*cpi-3B* locus (Ma et al. 2014). Fine mapping of this QTL shortened the genetic interval to 0.7 cM, containing 63 coding genes in the reference wheat genome (Zheng et al. 2015). Future map-based cloning and identification of the functional gene in this large-effect QTL may help elucidate the molecular bases of wheat resistance to FCR.

Other resistant QTLs have been identified using bi-parental populations. Early investigation discovered a resistant locus near the dwarfing gene *Rht1* on chromosome 4B from the wheat cultivar “Kukri” (Wallwork et al. 2004). Inherited from the wheat line “W21MMT70” with partial resistance to FCR, two QTLs were mapped to chromosomes 2D and 5D (Bovill et al. 2006). A major QTL on chromosome 1DL (*QCr*.*usq-1D1*) and several minor QTLs were identified in wheat line “2-49 (Gluyas Early/Gala)” using SSR markers (Collard et al. 2006; Collard et al. 2005). FCR resistance screening of 32 wheat genotypes identified “2-49”, “Aso zairai 11”, and “Ernie” as resistant sources. A QTL derived from “Ernie” was mapped to chromosome 3BL near markers *wPt-1151* and *wPt-1834* (Li et al. 2010). An Australian spring wheat cultivar “Sunco” showed partial resistance to FCR induced by *F. pseudograminearum*. Using bi-parental QTL mapping, a major QTL was identified on chromosome 3BL, between SSR markers *Xgwm247* and *Xgwm299* (Poole et al. 2012). These resistant sources of “W21MMT70”, “2-49”, and “Sunto” were then used for QTL pyramiding (Bovill et al. 2010). Four FCR-resistant QTLs were discovered, and their resistant alleles were derived from the bread wheat commercial variety “EGA Wylie”. Major QTLs on chromosomes 5DS and 2DL were consistently detected in all three populations and two minor QTLs were mapped to chromosome 4BS (Zheng et al. 2014). QTL mapping was also performed to find genetic loci controlling partial resistance to FCR in the four wheat germplasm “2-49”, “Sunco”, “IRN497”, and “CPI133817”. FCR resistance was evaluated in both seedlings and adult plants. Six QTLs among these resistant wheat genotypes were revealed (Martin et al. 2015).

A GWAS approach was used to screen 2,514 wheat genotypes for FCR resistance, and DArT and SSR markers identified two major QTLs on chromosome 3BL that explained 35% and 49% of the phenotypic variation (Liu et al. 2018). A set of 126 spring bread wheat lines from CIMMYT was phenotyped against FCR induced by *F. culmorum* and further genotyped using DArT markers, which resulted in the identification of three major QTLs on chromosomes 3B and 2D (Erginbasorakci et al. 2018). The use of GWAS for FCR resistance has greatly benefited from advanced high-throughput sequencing techniques and the released hexaploid wheat genome. A total of 234 Chinese wheat cultivars were evaluated for FCR resistance in four greenhouse experiments, with GWAS using a high-density 660K SNP assay. This revealed a major QTL on chromosome 6A, which was subsequently validated using a bi-parental population of “UC1110/PI610750” (Yang et al. 2019). The same team screened the FCR resistance of another 435 wheat introgression lines (generated by crossing of Yanzhan1 with other elite varieties) and performed GWAS using 660K SNP array. Most of the significant SNP associations were distributed on chromosome 4B and a gene encoding a dirigent protein (*TaDIR-B1*) was validated as a negative regulator of FCR resistance (Yang et al. 2021). A recent GWAS approach phenotyped 358 Chinese germplasm for FCR resistance, with less than 10% exhibing a lower disease index. The wheat 55K SNP assay was applied for association analysis, resulting in detection of significant QTLs on chromosomes 1BS, 1DS, 5DS, 5DL, and 7BL (Jin et al. 2020). GWAS was also performed to evaluate FCR resistance of 161 wheat accessions under growth room and greenhouse conditions using *F. culmorum* as the pathogen. Using a 90K SNP array, a total of fifteen QTLs for FCR resistance were detected with one major QTL on chromosome 3BS near the FHB resistance *Fhb1* locus (Pariyar et al. 2020). A marker-assisted recurrent selection approach was next performed on two populations to pyramid minor FCR-resistant QTLs. Using 9K SNP array, a total of 23 marker-trait associations were identified by GWAS (Rahman et al. 2020).

In **Supplementary Table 2**, we summarize wheat germplasm resistant to FCR induced by either *F. pseudograminearum* or *F. culmorum*. Identified QTLs controlling FCR resistance are also highlighted (**Supplementary Table 2**), with their genomic distributions annotated using the wheat genome database (**Fig. 3**).

**Fig. 3.**
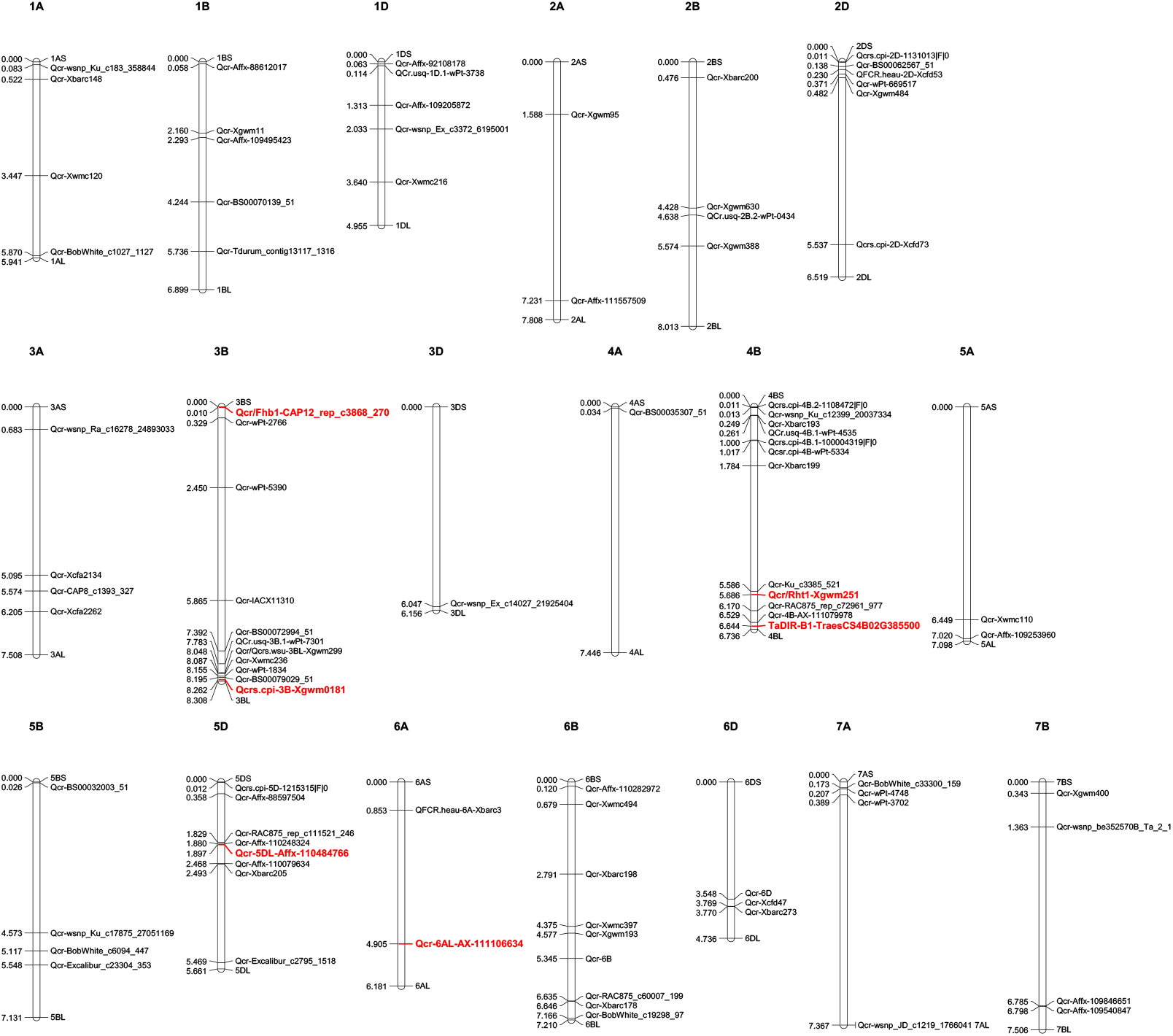
Genetic loci controlling wheat resistance to *Fusarium* crown rot. Molecular markers, SNPs, and genes associated with FCR-resistant QTLs were collected from previous publications and searched against the JBrowse-1.12.3-release of the common wheat “Chinese Spring” genome available from the “*Triticeae* Multi-omics Center (http://202.194.139.32/)”. Physical positions (numbers indicated on the left side of each chromosome) were used to generate a distribution map of all the collected QTLs using Mapchart v2.32 software. QTLs with major effect or linked with designated genes are highlighted in red. Detailed information for these QTLs can be found in **Supplementary Table 2**.

### Genetic determinants of wheat resistance to sharp eyespot

Wheat resistance to sharp eyespot is controlled by QTLs. However, additional efforts should focus on identification of resistant germplasm and genetic loci conferring resistance to this fungal disease. A recent large-scale screening of sharp eyespot resistant germplasm in Chinese wheat cultivars revealed no immune or highly resistant germplasm, and only 4% exhibiting moderate resistance to *R. cerealis* (Ren et al. 2020). Introgression of exogenous chromosome segments from wheat relatives might help generate novel resistant germplasms. For example, a wheat-rye 4R chromosome disomic addition line gained high resistance to sharp eyespot (An et al. 2019). Wheat cultivars “Luke” and “AQ24788-83” showed high resistance to *R. cerealis*. Subsequent genetic investigations revealed seven significant sharp eyespot resistant QTLs on chromosomes 1A, 2B, 3B, 4A, 5D, 6B, and 7B (Chen et al. 2013; Guo et al. 2017). Using 90 K SNP and SSR markers, five QTLs on chromosomes 2BS, 4BS, 5AL, and 5BS controlling resistance to *R. cerealis* were identified from the wheat cultivar “CI12633” (Wu et al. 2017). Three QTLs controlling resistance of wheat cultivars “Niavt14” and “Xuzhou25” to *R. cerealis* were mapped to chromosomes 2B and 7D (Jiang et al. 2016). A recent study using the same population of “Niavt14/Xuzhou25” and 55K SNPs revealed three novel stable QTLs on chromosomes 1D, 6D, and 7A (Liu et al. 2020).

In **Supplementary Table 3**, we summarize wheat germplasm resistant to *R. cerealis*. Reported QTLs controlling sharp eyespot resistance are highlighted (**Supplementary Table 3**), with their genomic distributions annotated using the wheat genome database (**Fig. 4**).

**Fig. 4.**
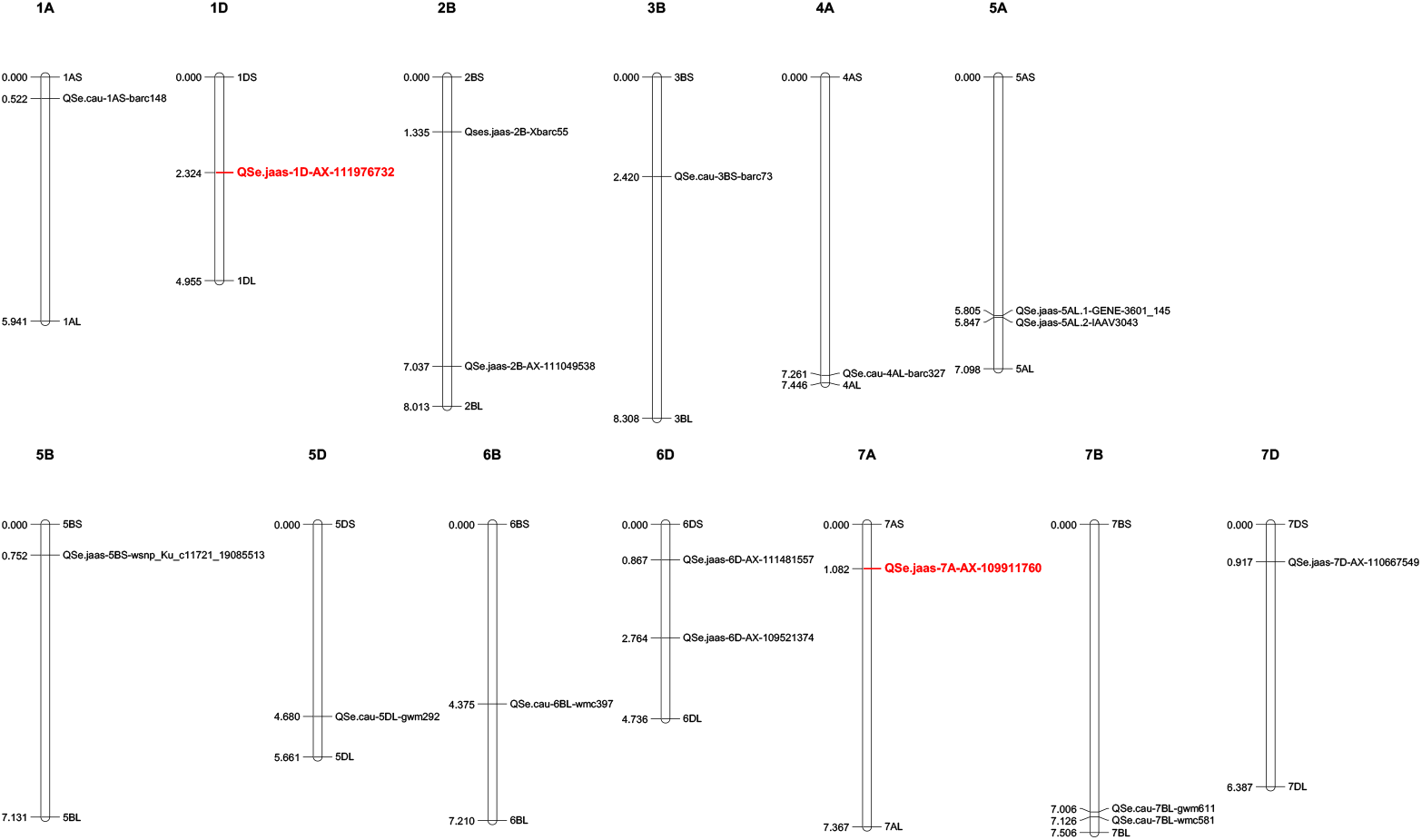
Identified QTLs controlling wheat resistance to sharp eyespot. Molecular markers associated with Rc-resistant QTLs were collected from previous publications and searched against the JBrowse-1.12.3-release of the common wheat “Chinese Spring” genome available from the “Triticeae Multi-omics Center (http://202.194.139.32/)”. Physical positions (numbers indicated on the left side of each chromosome) were used to generate a distribution map of all the collected QTLs using Mapchart v2.32 software. Detailed information for these QTLs can be found in **Supplementary Table 3**.

## Conclusion and future perspectives

We have described three rot diseases that commonly infect the stem base of wheat plants (**Fig. 1**). These diseases are major threats to wheat productions in wheat-maize rotation areas with large-scale application of straw returning. Wheat breeding is the most efficient way to control these devastating fungal diseases. However, as summarized in this review (**Supplementary Tables 1, 2, and 3**), there are few wheat germplasm with relative high resistance to *B. sorokiniana, F. pseudograminearum*, or *R. cerealis*. Large-scale screenings of resistant wheat germplasm are still urgently needed for effective wheat breeding applications. New germplasm resources including wheat relatives (*e*.*g*. introgression lines using *Thinopyrum ponticum, Triticum spelta*, and rye) may have great potential to improve wheat resistance to these root and crown rot fungal diseases.

Genetic improvement of wheat resistance to these diseases requires exploring novel QTLs that control resistance. There are several previously reported resistant QTLs (**Supplementary Tables 1, 2, and 3**) and their genomic distributions have been mapped based on the released wheat genome (**Figs. 2, 3, and 4**). Some identified QTLs that confer resistance to *B. sorokiniana* are associated with loci responsible for wheat resistance to other foliar fungal diseases, such as *Lr34/Yr18/Pm38, Lr46/Yr29*, and *Tsn1*. Wheat leaves might restrain the infection of different foliar fungal diseases using similar molecular approaches mediated by resistant genes. Wheat germplasm with broad-spectrum resistant loci should be evaluated for potential resistance to spot blotch or common root rot induced by *B. sorokiniana*. Of QTLs that control resistance to *Fusarium* crown rot, ones that also have resistance to FHB may be more valuable, since the major causal agents of these diseases (*F. pseudograminearum, F. culmorum*, and *F. graminearum*) are very likely to co-exist in a cultivation environment. For genetic studies on QTLs controlling resistance to sharp eyespot, the large-scale screening of resistant wheat germplasm would greatly accelerate the identification of novel QTLs correlated with resistance to sharp eyespot. There is also an urgent need to employ GWAS technique to screen for more sharp eyespot resistant QTLs at the genome-wide level.

Constructing near-isogenic lines and using residual heterozygotes allow the use of fine mapping and further positional cloning for key gene/loci that control resistance. With advanced genomic and capture-sequencing techniques such as MutRenSeq, AgRenSeq, and Exome Capture, fast-cloning approaches might accelerate this time-consuming process (Steuernagel et al. 2016; Arora et al. 2019; Krasileva et al. 2017). Gene editing may also increase the rate of genetic improvement of wheat resistance to these fungal diseases (Wang et al. 2018). Both forward and reverse genetic studies will provide valuable targets for the application of CRISPR-Cas9 in wheat.

Efforts should also be made to convert traditional markers used previously to identify resistant QTLs (microsatellite, SSR, and DrAT) to SNP markers, as SNP markers may serve as valuable tools for high-throughput marker-assisted selection in wheat breeding. Progress in wheat genome research and increased availability of high-density SNP toolkits will facilitate the use of GWAS on collected wheat germplasm to more efficiently identify novel resistant sources and genetic loci.

## Acknowledgements

The authors would like to thank Prof. Zaifeng Li from Hebei Agricultural University for discussion and input to this work.

## Declarations

### Authors’ contributions

Conceptualization, X.W., Z.K., and W.Z; Data collection, J.S., J.Z., S.Z., M.L., S.P.; Original draft preparation, X.W.; Review and editing, S.C. and F.C.; Supervision, X.W., Z.K., and W.Z. All authors have read and agreed to the published version of the manuscript.

### Data availability

Not applicable.

### Ethics approval and consent to participate

Not applicable.

### Consent for publication

Not applicable.

## Figure legends

**Supplementary Table 1.**
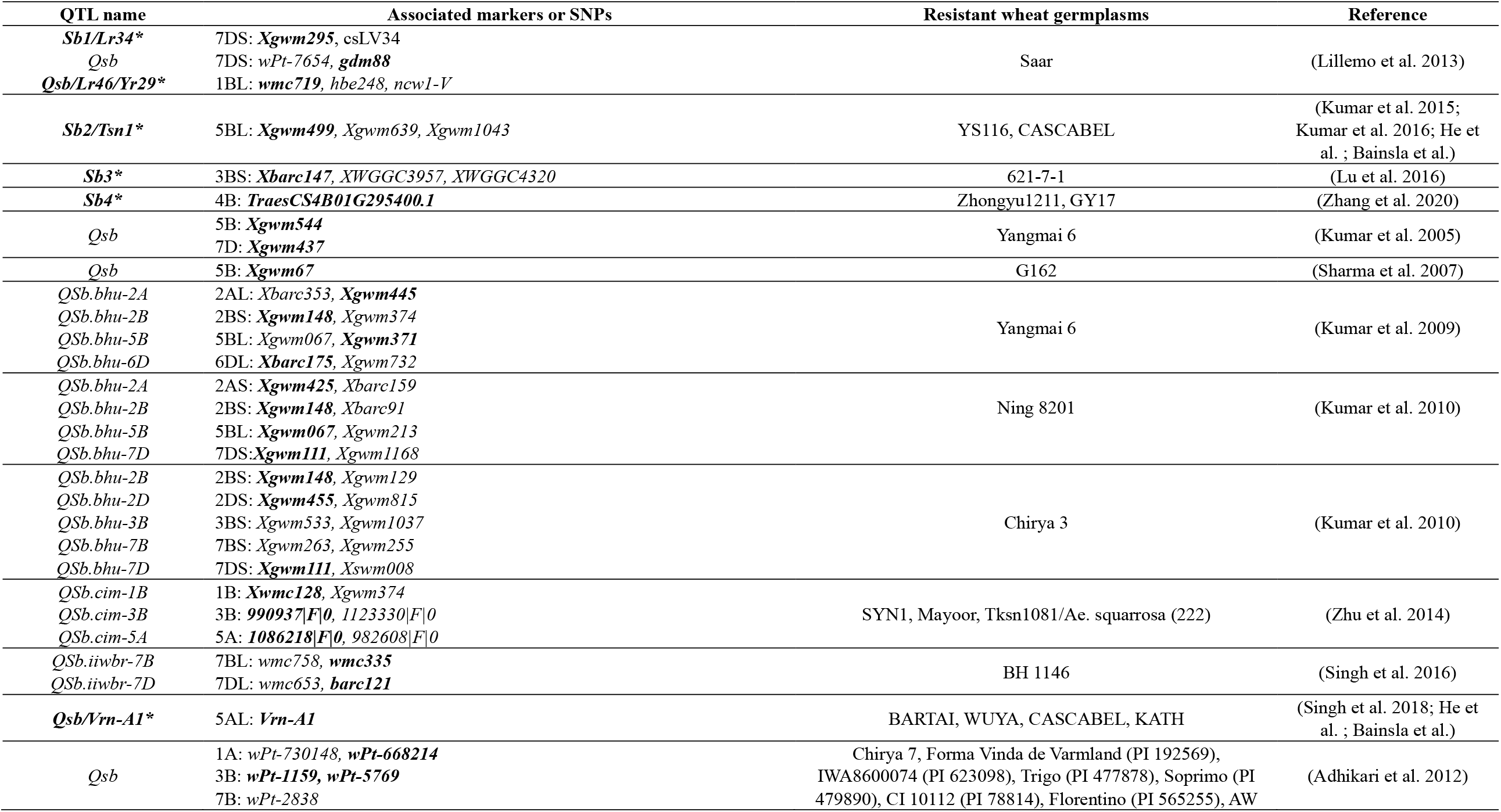
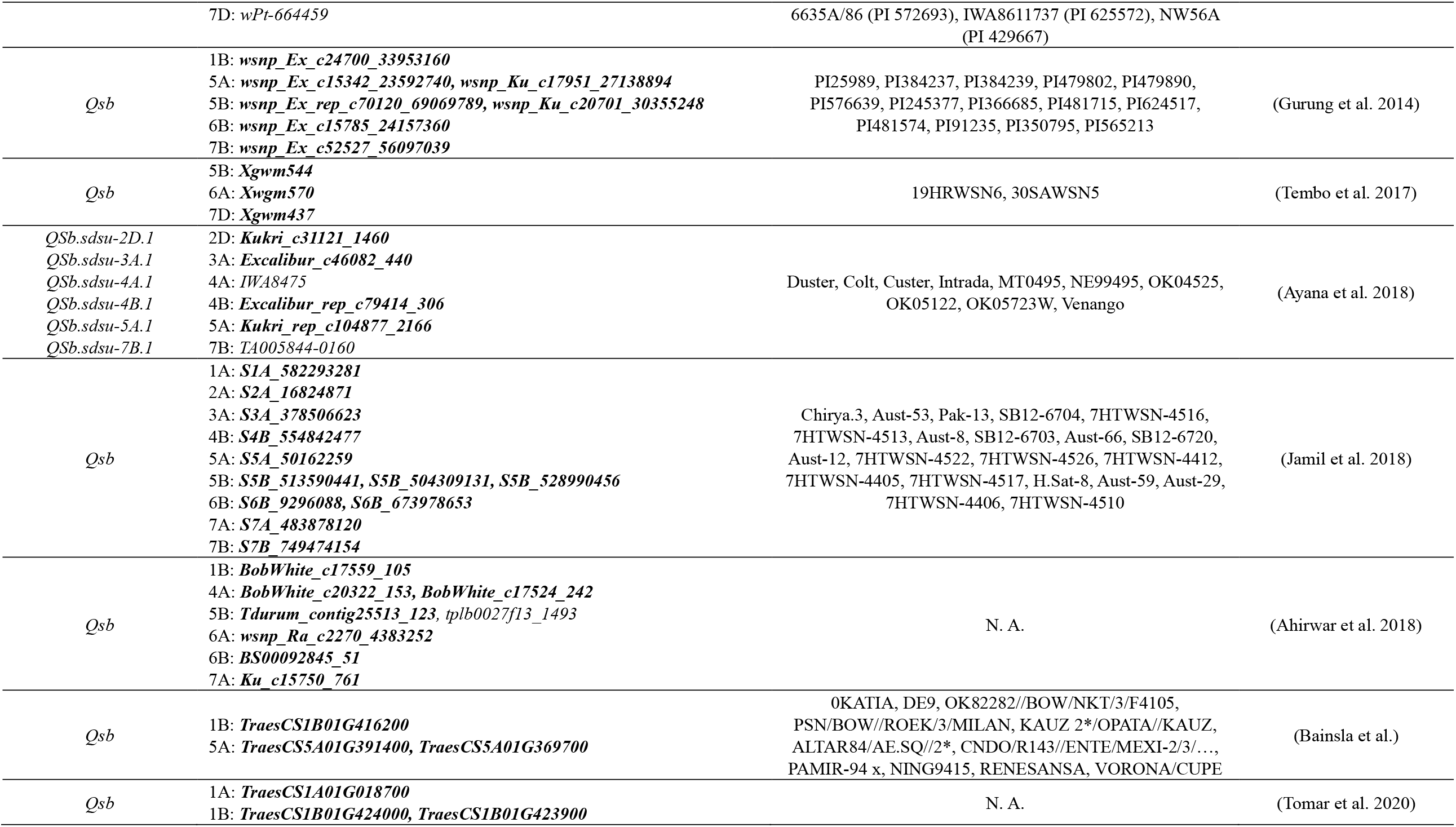
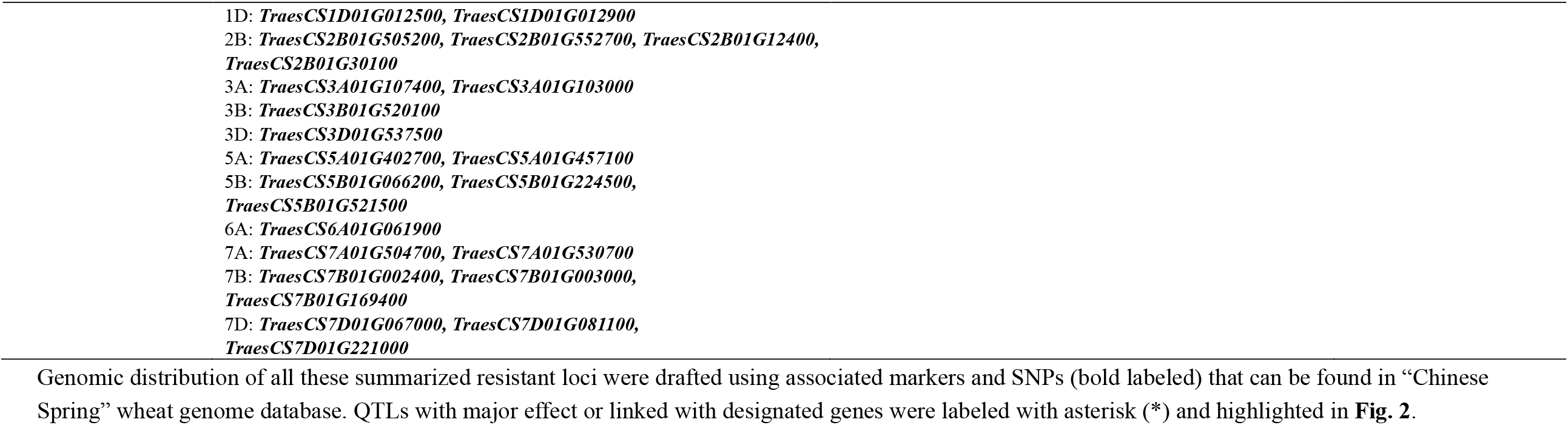
Genetics of resistance to common root rot (spot blotch) in wheat.

**Supplementary Table 2.**
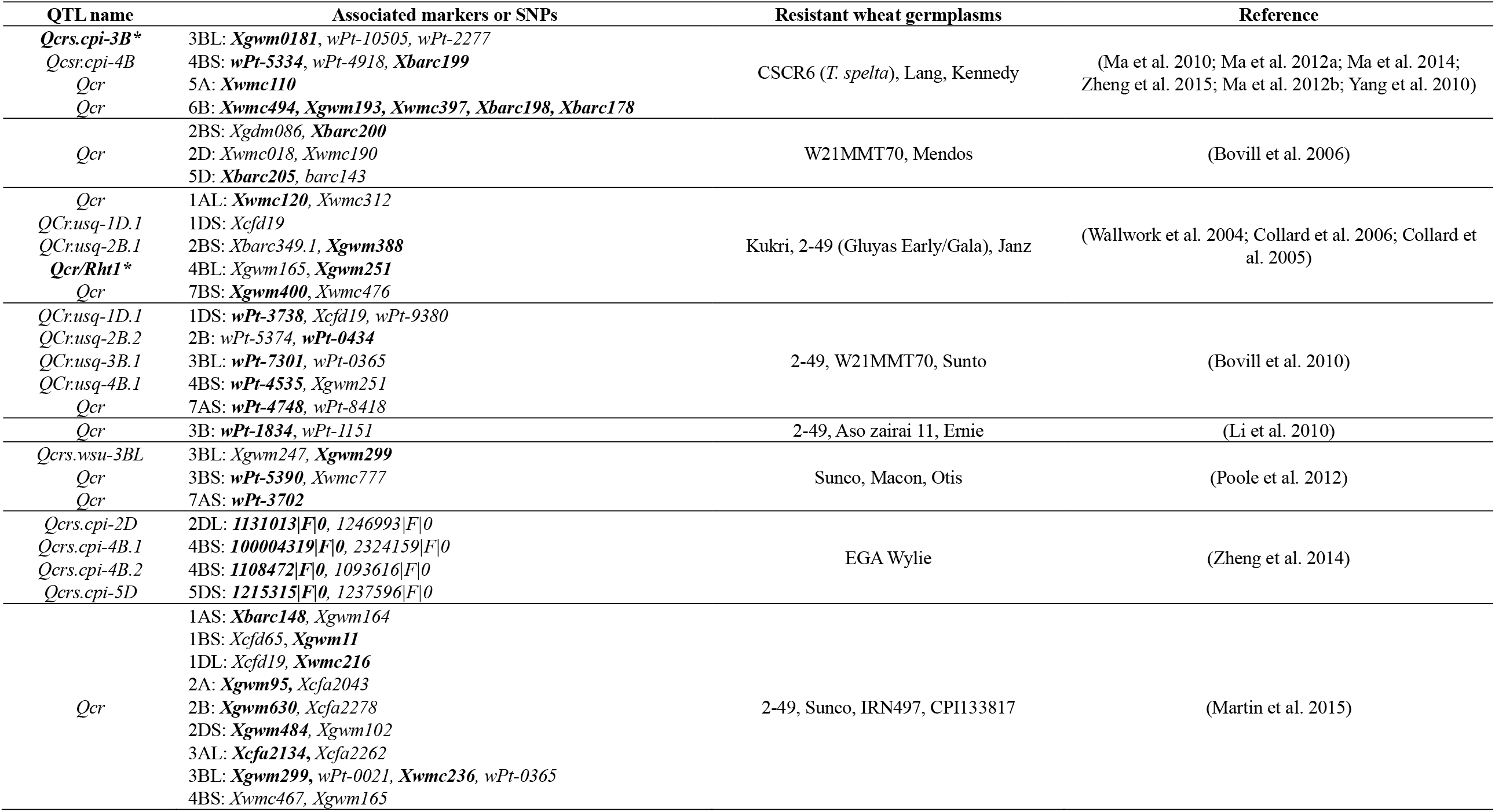
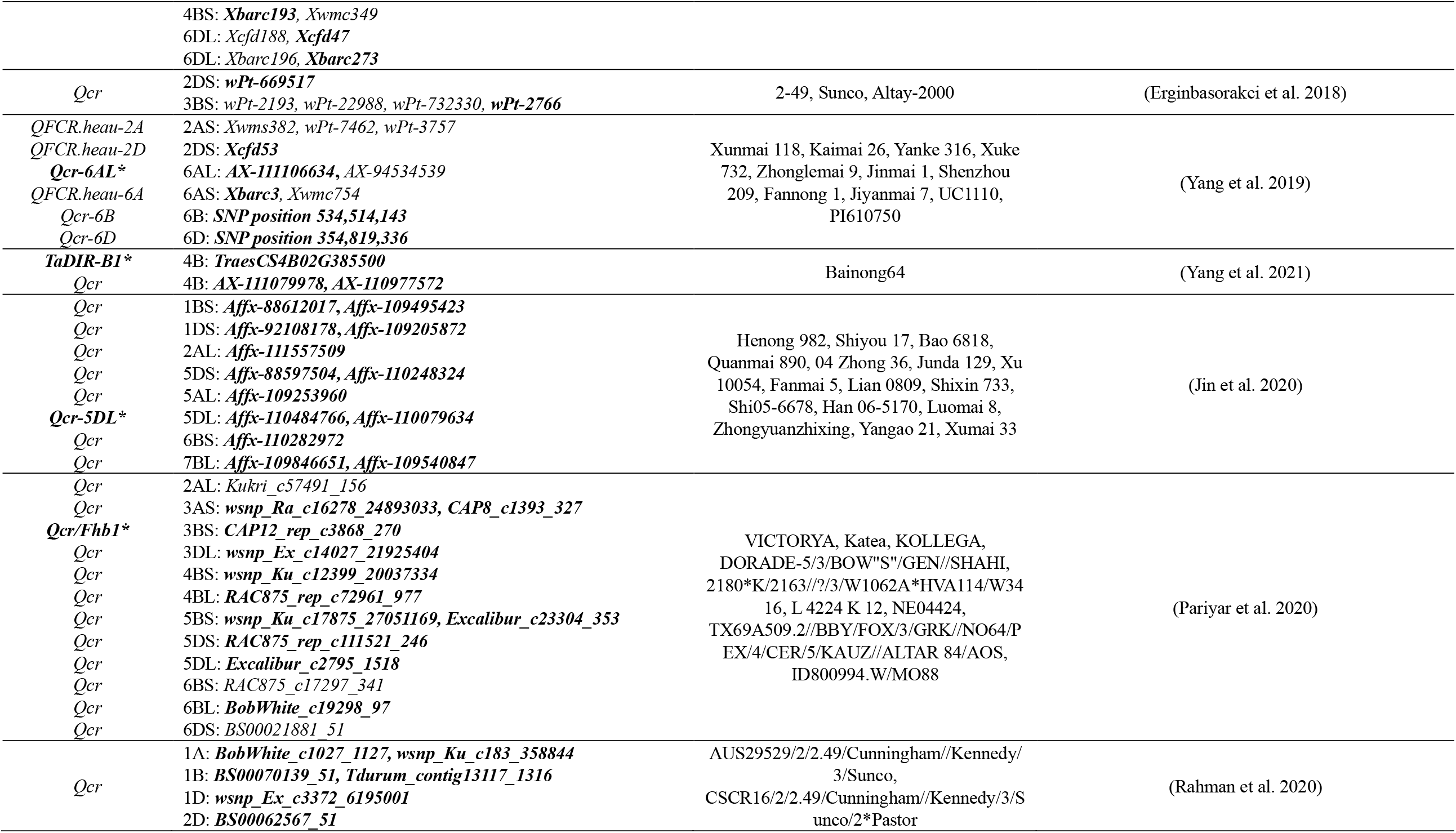
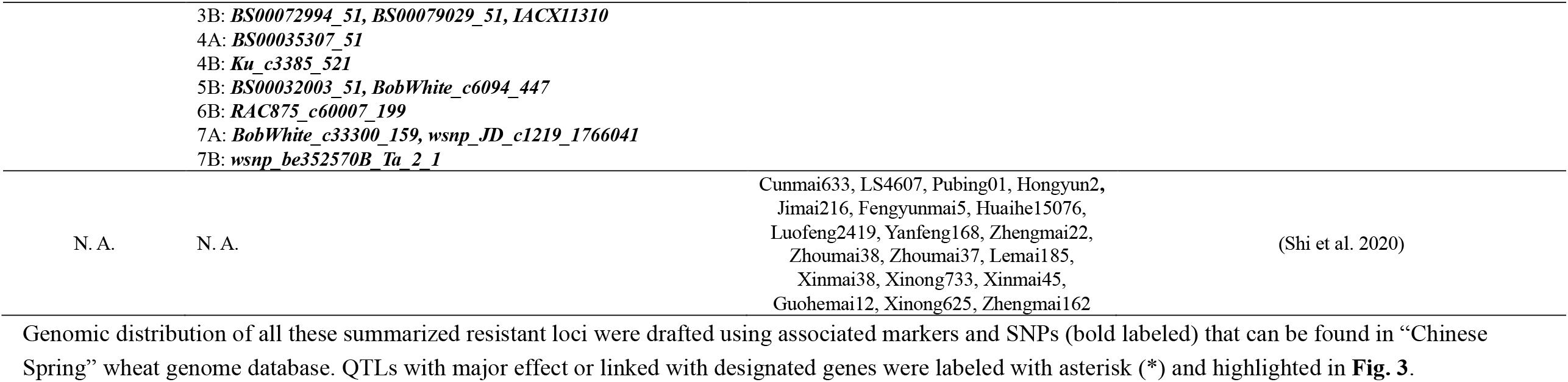
Genetic loci controlling wheat resistance to *Fusarium* crown rot.

**Supplementary Table 3.**
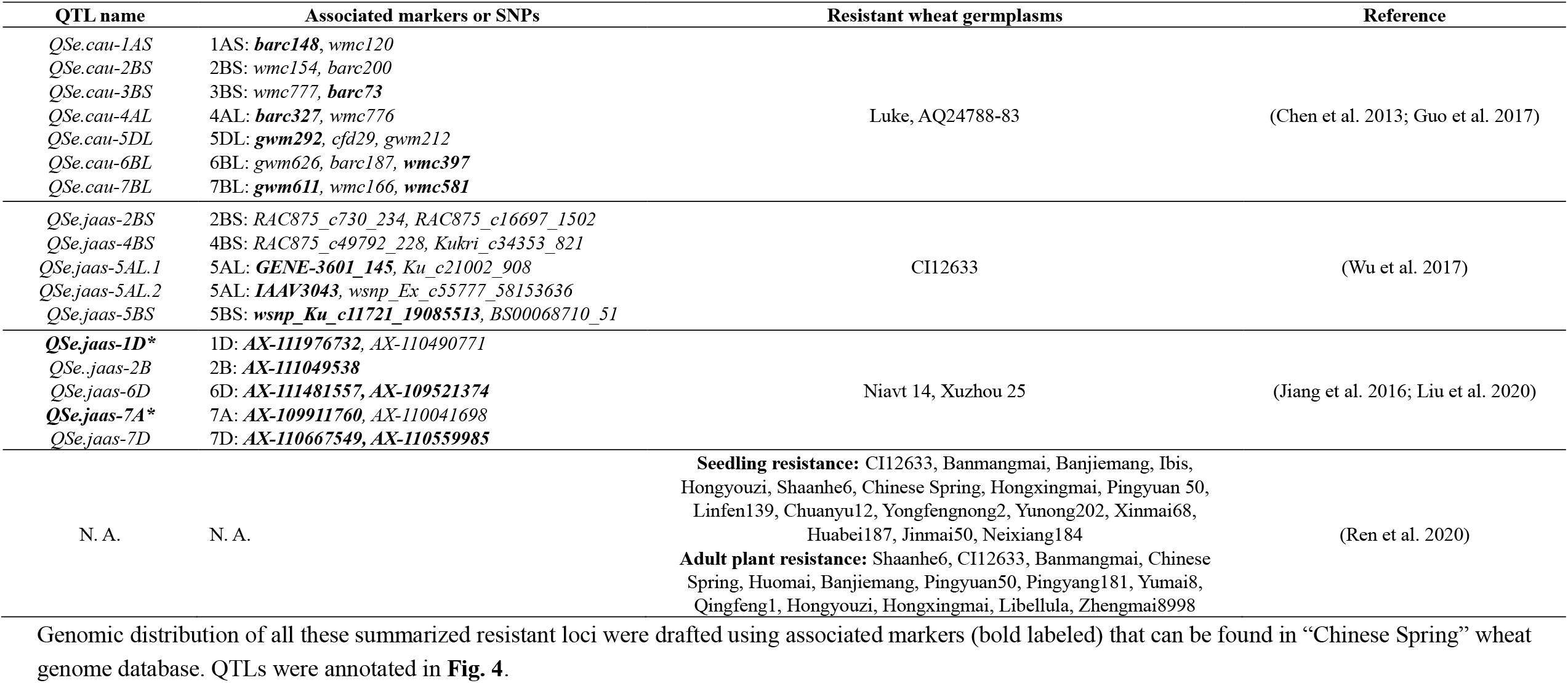
Genetic determinants of wheat resistance to sharp eyespot.

